# Drivers of plant intraspecific variation are trait-specific

**DOI:** 10.1101/2022.12.29.521136

**Authors:** Jianhong Zhou, Ellen Cieraad, Peter M. van Bodegom

## Abstract

- The importance of intraspecific trait variation (ITV) in community dynamics is increasingly being recognised, but the drivers of ITV have not yet been well-studied. Here, we analysed whether environmental conditions, biotic interactions and species features are related to ITV on a global scale.
- We compiled a global species’ ITV database including 2064 species which occurred in 1068 communities (plots) across 19 countries with 11 functional traits. The magnitudes of species’ ITV in this database were calculated according to the trait-gradient analysis which is independent of the length of the environmental gradient.
- We found that different traits had different main drivers, so we consider the drivers of ITV to be trait-specific. However, our findings still brought some order among these idiosyncratic patterns: leaf economics spectrum traits were more related to environmental conditions and leaf morphology traits were more related to biotic interactions. Size-related traits were related to both abiotic and biotic conditions.
- Our research suggests that the drivers of ITV deviated from the drivers of the mean trait values and that some trait coordination may fall apart upon climate change. Thus, our analysis enhances our understanding of trait variation and has important implications for models predicting vegetation responses to environmental change.

## 1 Introduction

The interrelationships between biodiversity, ecosystem functioning and ecosystem services form the basis of our understanding of ecosystem dynamics (van Bodegom & Price, 2015). Functional diversity, defined through the trait variation of the organisms, is an important facet of biodiversity and has increasingly been investigated in recent decades to better understand ecological dynamics. Fukami *et al*. (2005) provided the first empirical proof that trait assembly is convergent while it is idiosyncratic for species richness in the same community in a 9-years experiment. This implies that we can predict the trait assembly leading to functional diversity once we know the environmental drivers causing convergence (van Bodegom *et al*., 2014). Since then, traits are increasingly being used and quantified to understand the interactions between species and their environment.

With increasing knowledge of trait dynamics, it has become clear that traits do not only vary between species but also within species. Therefore, there has been a growing interest in incorporating intraspecific trait variation (ITV) into assembly theory and models to explain and predict trends in plant response to certain environmental conditions (McGill *et al*., 2006; Körner, 2018). Various studies have shown that, at both local and global scales, the intraspecific variation can be substantial, and can even equal the magnitude of interspecific variation (Messier *et al*., 2010; Kichenin *et al*., 2013; des Roches *et al*., 2018). This indicates that intraspecific trait diversity plays a pivotal role in community dynamics and ecosystem functioning (Bolnick *et al*., 2011). Albert *et al*. (2010) advanced our knowledge about ITV by quantifying the extent, structure and source of the intraspecific variability of plant traits (maximum height, leaf dry matter and leaf nitrogen content) in communities located in the central French Alps across a large climatic gradient. They also suggested that further research was required to compare interspecific and intraspecific variation. Shortly after, Lepš *et al*. (2011) proposed a method that could disentangle the role of interspecific and intraspecific trait variability to community variability. Their case study showed that all effects of intraspecific trait variation, interspecific trait variation and their covariation need to be assessed to understand the response of trait composition to the environment. Siefert *et al*. (2015) and two recent global meta-analyses (des Roches *et al*., 2018; Raffard *et al*., 2019) further showed the relative extent of intraspecific compared to interspecific trait variation within and among communities.

These studies show the critical importance of ITV for community dynamics (Westerband *et al*., 2021). However, while more and more studies have demonstrated that ITV may significantly structure communities and shape ecosystems, studies investigating the drivers of ITV are still lacking. Based on general ecological concepts, we may assume that species’ generic features as well as their strategies to deal with environmental heterogeneity and biotic interactions may influence ITV.

In an experiment with species of two different growth forms in German grasslands, Herz *et al*. (2017) showed that the ITV of grasses was much higher than that of forbs for almost all traits while Siebenkäs *et al*. (2015) found that only aboveground traits of grass species were more plastic than that of forbs in a common garden experiment. In contrast, (Albert *et al*., 2010b) did not find clear patterns between growth form and ITV. In a Chilean temperate rainforest, Fajardo & Siefert (2019) tested if species with a wider niche breadth had higher ITV. They found that ITVs of leaf nitrogen and phosphorus content were positively related to the soil phosphorus niche breadth, while the ITV of leaf area was negatively related to the light niche breadth. ITV has also been considered to link with plant strategy in terms of resource acquisition (Sultan & Bazzaz, 1993; Valladares *et al*., 2000). Grime (1977) defined three primary strategies for vascular plants regarding their response to stress and disturbance. He suggested that competitive plants (C-strategy) would have a higher ITV in photosynthesis-related characteristics than stress-tolerant plants (S-strategy). A decade later, a such pattern was indeed observed by Crick & Grime (1987) for two herbaceous species with these two contrasting strategies. Grassein *et al*. (2010) compared the ITV between two subalpine species and found that, compared with the conservative one, the exploitative species expressed an overall higher level of ITV in three functional traits (specific leaf area, leaf dry matter content and leaf nitrogen). Biotic interactions have also been suggested to drive ITV differences among species. Abakumova *et al*. (2016) found in common garden experiments that those species which encountered a high frequency of neighbouring individuals had more plastic morphological traits.

Despite the wealth of examples evaluating individual drivers of intraspecific trait variation, all experimental studies so far were conducted with few species or in a specific location. The only global-scale research (Stotz *et al*., 2021), to our knowledge, about the predictors of ITV across a large number of species (272 species), found that the drivers of plant phenotypic plasticity differed between trait categories. However, their research only focused on one component of ITV: phenotypic plasticity, whereas ITV is likely a combination of phenotypic plasticity and genetic differences among different populations of a species. Moreover, this study mainly assessed the influence of different environmental conditions (latitude, climatic variation and water/nutrient conditions) on trait plasticity, while we know that other drivers, including biotic interactions and species features that allow species to cope with abiotic/biotic variations (e.g. growth form, CSR strategy) may also affect ITV.

Therefore, our study aimed to comprehensively evaluate the relative importance of environmental conditions, biotic interactions and species’ features for intraspecific trait variation globally. To achieve this, we compiled a global database of intraspecific trait variation and species characteristics across 2064 species for 11 traits. This database allows us to test the main research questions: (1) Do environmental conditions, biotic interactions and species’ features relate to ITV on a global scale? (2) Does the ITV of traits related to similar trait groups (i.e. traits representing leaf economics, size or morphology) have similar drivers?

## 2 Materials and methods

### 2.1 Defining intraspecific trait variation based on trait-gradient analysis

We defined and determined estimates of ITV using the trait-gradient analysis, as outlined by Ackerly & Cornwell (2007). This analysis allows estimating ITV normalizing for the length of the environmental gradient involved. Compared to other metrics of intraspecific trait variation that most commonly express intraspecific variation as a percentage of the mean trait value of a given species (Albert *et al*., 2010b; Messier *et al*., 2010; Violle *et al*., 2012), the estimates of ITV by the trait-gradient analysis are much less affected by the length of the environmental gradient for which observations of an individual species are available (i.e. the range over which a trait of a species varies is likely to increase with an increasing length of an environmental gradient). At the same time, like other metrics, our ITV metric is unitless and thus allows direct comparison across traits and species.

At the plot level, the mean trait value (which is used to calculate ITV) can be mathematically expressed as:

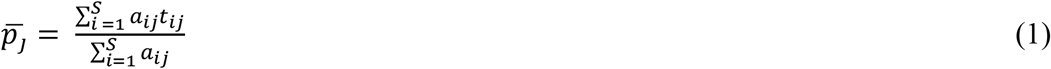

Where 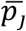, plot mean trait value, is defined as the abundance-weighted mean trait values of all the species in plot *j*. Given that traits may be considered to converge under the influence of environmental pressures, 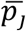 represents the position of a plot along the environmental gradient driving this trait (Ackerly & Cornwell, 2007). *t*_*ij*_ is the individual species trait value of a species *i* in plot *j, a*_*ij*_ is the abundance of a species *i* in plot *j*, and the total number of species in plot *j* is S.

ITV is expressed as relative to the (community-weighted) trait variation in the community. If one visualizes the variation of individual species trait values *t*_*ij*_ vs. the plot mean trait values 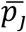 (Fig. **1**), sets of points (grey dots) align vertically at a particular value of 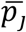, which indicates the species that occur together in the same plot j. A weighted least squares (WLS) regression through all *t*_*ij*_ *vs*. 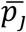 represents the community trait variation which, by the definition of 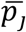, falls on a 1:1 line (represented in Fig. **1** by the dashed line). For an individual species, the slope of the WLS regression line of *t*_*ij*_ *vs*. 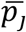 for that species, reflects the magnitude of ITV of a species. An example for the species *Embothrium coccineum* J.R.Forst. & G.Forst with 124 observations in our global database is shown in Fig. **1**.

**Fig. 1.**
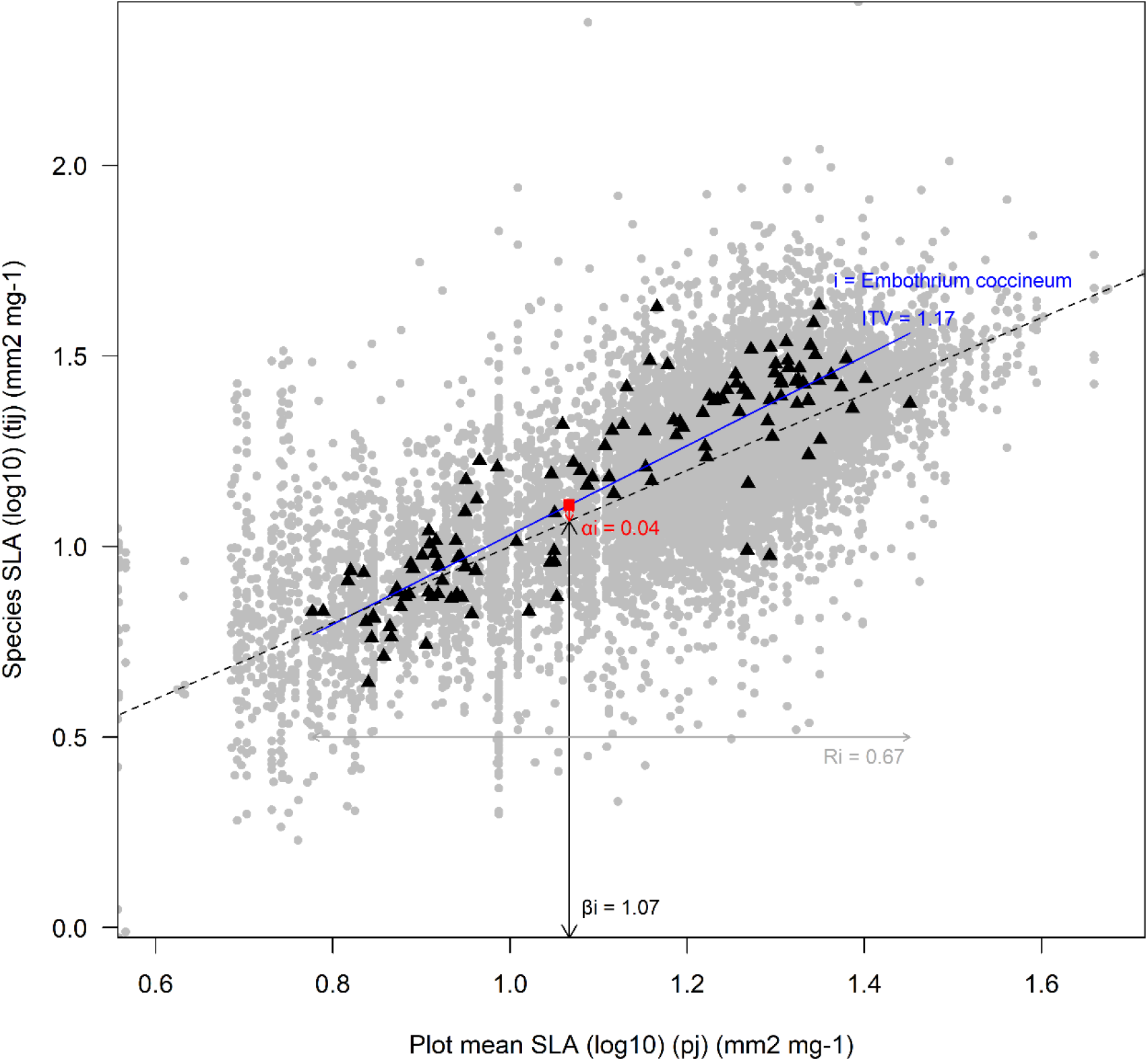
Scatterplot of individual species trait values (t_ij_) vs. plot mean trait values 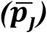 for log_10_-transformed specific leaf area (SLA) (mm^2^ mg^-1^) in our trait database. The dashed line is the 1:1 line. Black triangles illustrate the observations of Embothrium coccineum. The blue line is the weighted ordinary least squares (WLS) regression line for this species and the slope of this line reflects the intraspecific trait variation (ITV) of Embothrium coccineum. The red square shows its species mean trait value of log_10_-transformed SLA 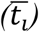 at its species mean niche position (β_i_) along the environmental gradient expressed by log_10_-transformed SLA. Species alpha niche position (α_i_), the difference between 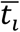 and β_i_, or the distance of the red square from the 1:1 line (marked as the red segment with arrows here). The grey segment with arrowheads represents the niche breadth of Embothrium coccineum along this trait gradient. See text for further explanation of these parameters.

An added value of the trait-gradient analysis is that, in addition to calculating ITV, it also allows determining additional species characteristics that act as indicators for some of the drivers we aimed to test (see section 2.3). The species mean trait value,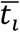:

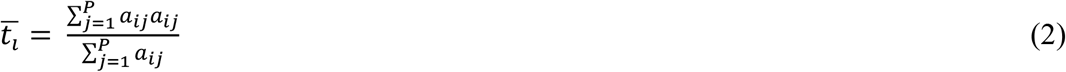

is defined as the abundance-weighted mean trait values of a species *i* across all plots, with *P* being the total number of plots where this species occurs. This allows calculating species beta niche position, *β*_*i*_:

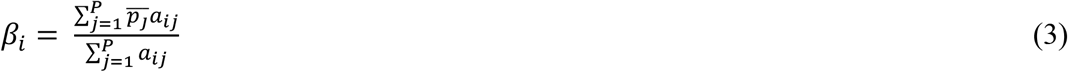

which is defined as the abundance-weighted mean of 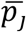 where species *i* occurs. *βi* represents the species mean niche position along the trait gradient. So, we regard *βi* as an indicator of the preference of a species across distinct environmental conditions.

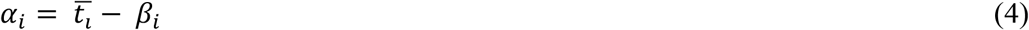

*αi*, species alpha niche position, is defined as the difference between the species mean trait value 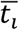 and the species beta niche position *βi. α*_*i*_ represents the trait values of a species *i* relative to that of co-occurring species. Because *αi* differentiates a species *i* from co-occurring species, we consider *αi* to be an indicator of the biotic interactions.

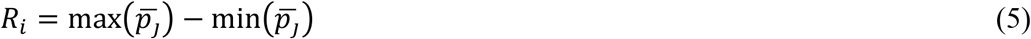

*R*_*i*_, species niche breadth, is defined as the range of 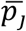, where a species *i* occupies along the trait gradient.

The example species, *Embothrium coccineum, with these values* is illustrated in Fig. **1**. The red square 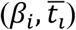 shows its species mean niche position (*β*_*i*_) and its species mean trait value 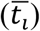. Its *β*_*i*_ = 1.07 (log_10_ mm^2^ mg^-1^), close to the median of the trait gradient, indicates that this species tends to occupy median SLA plots. Its *α*_*i*_ = 0.04 (log_10_ mm^2^ mg^-1^), representing the species mean trait value 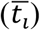 of *Embothrium coccineum* is 0.04 (log_10_ mm^2^ mg^-1^) higher than the corresponding plot mean trait value 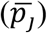 where *β*_*i*_ is, indicates that *Embothrium coccineum* has slightly higher SLA values compared to that of co-occurring species. The niche breadth *R*_*i*_ of *Embothrium coccineum* along the trait gradient equals 0.67 (log_10_ mm^2^ mg^-1^).

We repeated the trait-gradient analysis procedure to calculate ITVs and above variables for all 11 traits. For most traits, except for LDMC, LCC and SSD, their original trait values did not comply with normal distribution, thus their log_10_-transformed trait values were used in these calculations.

### 2.2 Database preparation

The database used in this study was developed from the species’ ITV database compiled from 19 different studies (see detailed reference list in Table S1 of Zhou *et al*. 2022)). In brief, this database contains ITV estimates for 2064 vascular species and 11 functional traits, namely specific leaf area (SLA), leaf size (LS), leaf dry matter content (LDMC), leaf nitrogen content (LNC), maximum height (MH), leaf phosphorus content (LPC), leaf carbon content (LCC), leaf thickness (LT), leaf tissue density (LTD), stem specific density (SSD), and specific root length (SRL). Species observations covered tropical, temperate and boreal biomes (Zhou *et al*., 2022).

For the current study, we expanded the database containing the magnitude of ITV for each species (Zhou *et al*., 2022) with the calculation of three additional indicators of species characteristics (species alpha niche position, species beta niche position and species niche breadth; see section 2.1) using the same trait-gradient analysis (Ackerly & Cornwell, 2007) and we added two species features (growth form and CSR strategy).

For plant growth form, we amalgamated growth form data from four public Categorical Plant Traits Databases from the TRY database (Kattge *et al*., 2011; dataset IDs: 57, 104, 110, 117; https://www.try-db.org/de/Datasets.php) after updating their species names using the R package *Taxonstand* (Cayuela *et al*., 2012).We categorized species according to six growth forms: ferns and allies, lianas, herbs, graminoids (sedges, rushes and grasses), shrubs and trees. If a growth form category for a species was inconsistent across the databases or was not available, we performed an online search for information on these species and one by one completed the growth form values for all species in our database.

To assign species with CSR strategy values, we used a combination of calculated and literature values. For those species with trait values of LS, LDMC and SLA, we calculated CSR strategy values according to the method of Pierce *et al*. (2017). We also added CSR values from the supplementary Table1 of Pierce *et al*. (2017) and a CSR-lookup table for British plant species (Hill 2004). For any species with multiple CSR values available, we selected the common denominator. Finally, we transformed CSR values into the percentage of C, S, and R.

Finally, we considered that the species niche breadth estimates are more sensitive to the number of observations (the number of plots) than ITV itself. Therefore, we only tested ITV against species niche breadth for those estimates that were based on a sufficient number of observations, as the species niche breadth calculated from a sufficient number of observations becomes stable and reliable. To determine the threshold number of observations for calculating species niche breadth, we took the species with the most observations (134 for SLA) in the database, *Amomyrtus luma* as an example. First, we bootstrapped an increasing number of observations, calculating species niche breadth for each draw. We found that species niche breadth fitted a saturation curve *vs*. the number of observations (see Supplementary Information Fig. **S1**). Subsequently, we tested how many observations the calculated species niche breadth did not differ significantly from that calculated for 134 observations. This minimum number of observations was 32 for SLA, coinciding with around 85% of the maximum niche breadth. This number of observations was subsequently taken as the threshold number for providing reliable estimates for species niche breadth, assuming that the same threshold applied to other traits.

### 2.3 Framework and metrics

**Fig. 2.**
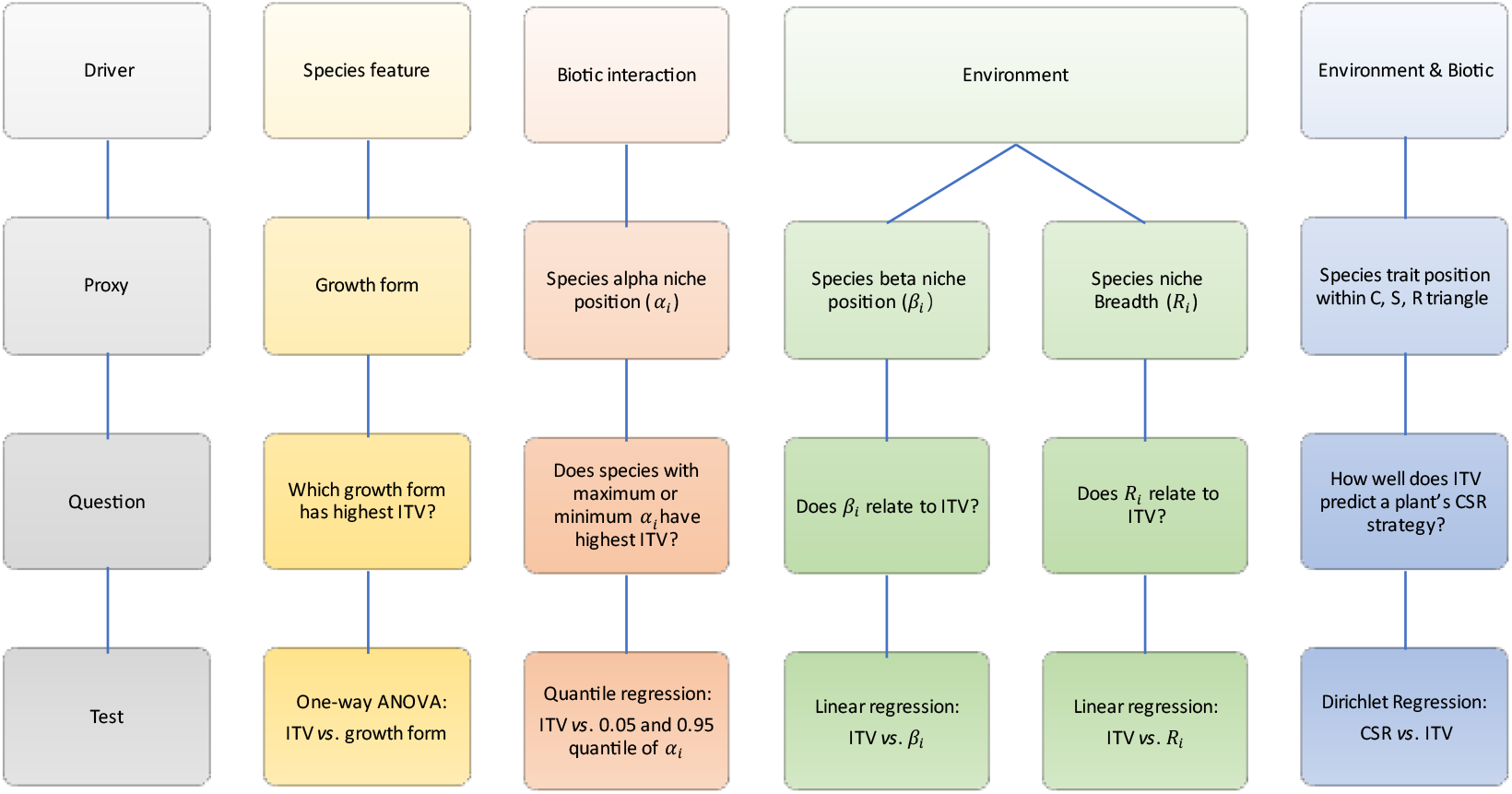
Overview of our framework and the metrics used to test for the global drivers of intraspecific trait variation (ITV) for this study. We aimed to test whether environmental conditions, biotic interactions and species’ features are generic drivers of ITV, using the framework outlined in Fig. **2**. Growth form is used as a generic indicator of species appearance. For biotic interaction, we used alpha niche position () as a proxy for biotic interactions, as it refers to the variation of a species’ trait relative to other species in the same community (Ackerly & Cornwell, 2007). For environmental conditions, as our database does not contain climatic nor soil conditions for all observations, we used species beta niche position () and species niche breadth (R_i_) as proxies. As indicated by Ackerly & Cornwell (2007), the beta niche position refers to the observed species position across geographic gradients and the species niche breadth refers to the range that species occupies along environmental gradients. Finally, the CSR strategy reflects the strategy of a species in relation to environmental and biotic stress.

### 2.4 Statistical analysis

We used multiple statistical methods to test the relationships between our drivers and ITV (Fig. **2**). Each test was run separately for each trait analysed. We used a one-way ANOVA to test if growth form affected ITV, using a Tukey post-hoc test to assess which growth forms differed significantly from each other (for a given trait). Based on the ecological concept of limiting similarity, we assumed that the maximum or minimum species alpha niche position (*α*_*i*_) would have the highest impact on ITV, so we used quantile regression to test if the species with maximum or minimum *α*_*i*_ had the highest ITV. To evaluate whether and how environmental conditions were related to ITV, we applied linear regressions to test the relationship between ITV and the species beta niche position *α*_*i*_ and the species niche breadth *R*_*i*_. Lastly, we used Dirichlet regression (Douma & Weedon, 2019) to test whether ITV is related to CSR strategies. Given that the CSR strategy for each species was described by three variables (percentages of C, S and R) that co-vary and sum to one, Dirichlet regression analysis was chosen here. The distribution of species trait positions within the C, S and R triangle in the database is shown in Fig. **S2**. All analyses were conducted in R (version 3.6.3, R Core Team (2020)) and Microsoft Excel 2019.

## 3. Results

ITV varied strongly between species. The ITV values of different traits for most species were mostly distributed between 0∼1 (Fig. **S3**), indicating that the trait variation of most species is lower than the change in the community trait variation (which has the value 1 by definition, including the trait variation induced by ITV and species turnover) with changing environmental conditions. Some species had greater ITVs than that of the community (ITV > 1), while a few species had ITVs < 0, the latter implies counter-gradient variation (the direction of the trait change for these species was opposite to the trait variation direction of their community).

ITV did not vary strongly among different traits (R2 = 0.038). Based on the Tukey post-hoc tests for the ITV of different traits (Fig. **S4**), the mean ITV was highest for SRL (albeit with a relatively small sample size), followed by LPC, MH and LCC. SLA, LT, LNC, LTD, SSD and LDMC were less plastic, and LS was the most stable trait.

### 3.1 General overview of drivers of intraspecific trait variation across traits

Overall, all the five drivers (growth form, species alpha niche position, species beta niche position, species niche breadth and CSR strategy) were related to ITV (Table **1**), but R^2^ associated with the tests were mostly lower than 10% (except for the effects of growth form and species beta niche position to LCC, of niche breadth and species alpha niche position to LDMC, and species alpha niche position to leaf thickness). This means that none of the individual drivers was very strongly related to the ITV of most traits and that major sources of variation in ITV remained unexplained. Moreover, for all traits, their ITV was found to be significantly related to at least one driver but none of the drivers was significantly related to all traits. Note that the number of observations varies for different tests, with fewer observations for *R*_*i*_ and CSR strategies than that for the other three proxies. This is caused by the lack of data on CSR strategy for some species and the strict threshold we had set for the minimum number of observations to calculate a reliable species niche breadth. The results for each driver are discussed in the next sections.

**Table 1.**
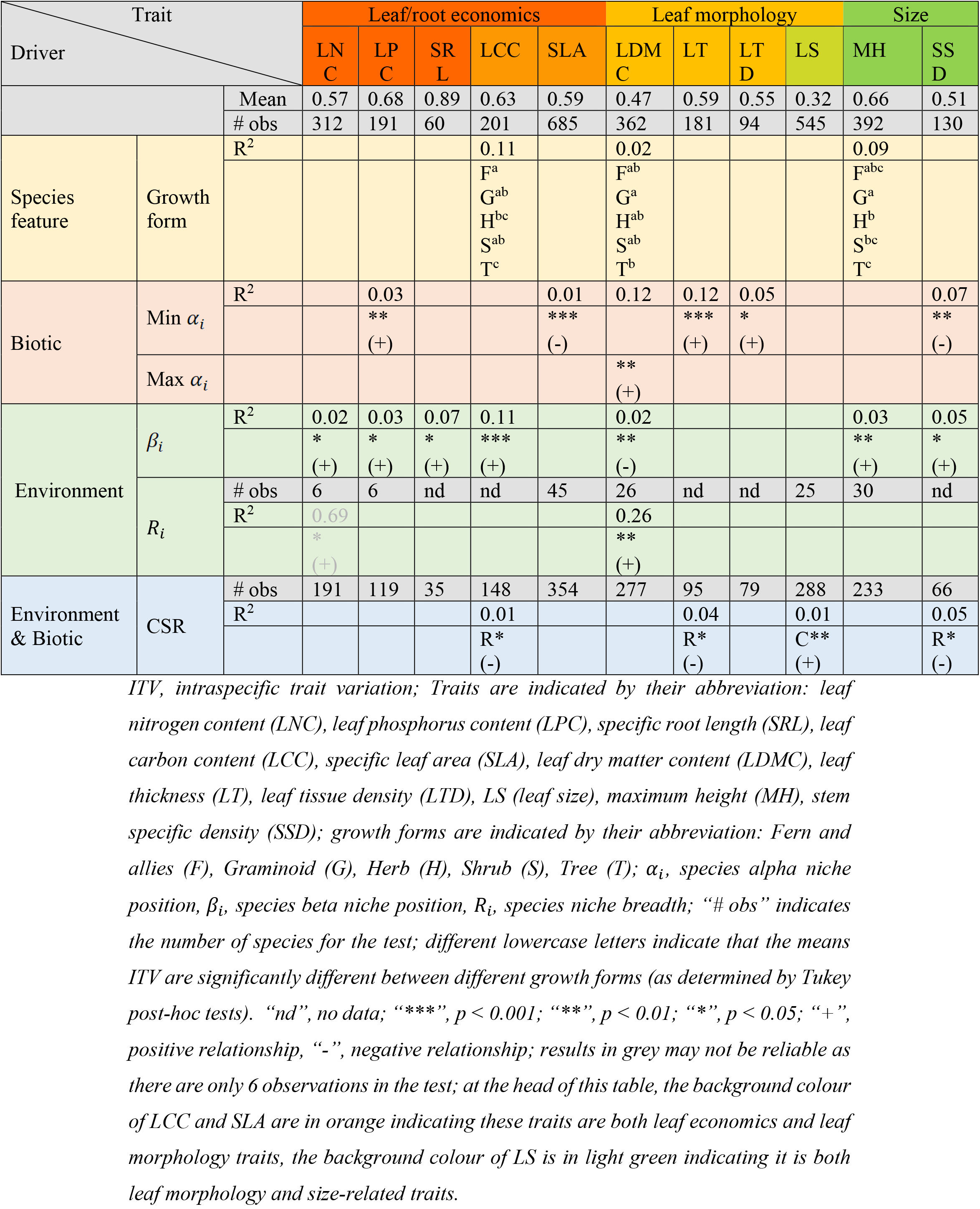
Overview of the results of statistic tests on the drivers of the intraspecific trait variation (ITV) of different traits.

### 3.2 Intraspecific trait variation and growth form

The ITVs of three traits (LDMC, MH and LCC) were related to growth form (Fig. **3**). For LDMC, the ITV of graminoids was significantly different from that of trees (Fig. **3B**, R^2^ = 0.02). For MH, trees had the highest ITV while graminoids had the lowest ITV (Fig. **3E**, R^2^ = 0.088). The ITV of trees was also significantly higher than that of herbs (p < 0.01), but not significantly different from that of shrubs and fern and allies. The ITV of shrubs (p < 0.05) and of herbs (p < 0.01) was significantly higher than that of graminoids. For LCC, trees had the highest ITV while ferns and allies had the lowest ITV (Fig. **3H**, R^2^ = 0.115, but only 4 observations for ferns and allies). The ITV of trees was significantly higher than that of graminoids (p < 0.05), shrubs (p < 0.01) and ferns and allies (p < 0.05) but not significantly different from that of herbs. Herbs also had higher ITV than that of ferns and allies (p < 0.05). There were no significant patterns for the other eight traits.

**Fig. 3.**
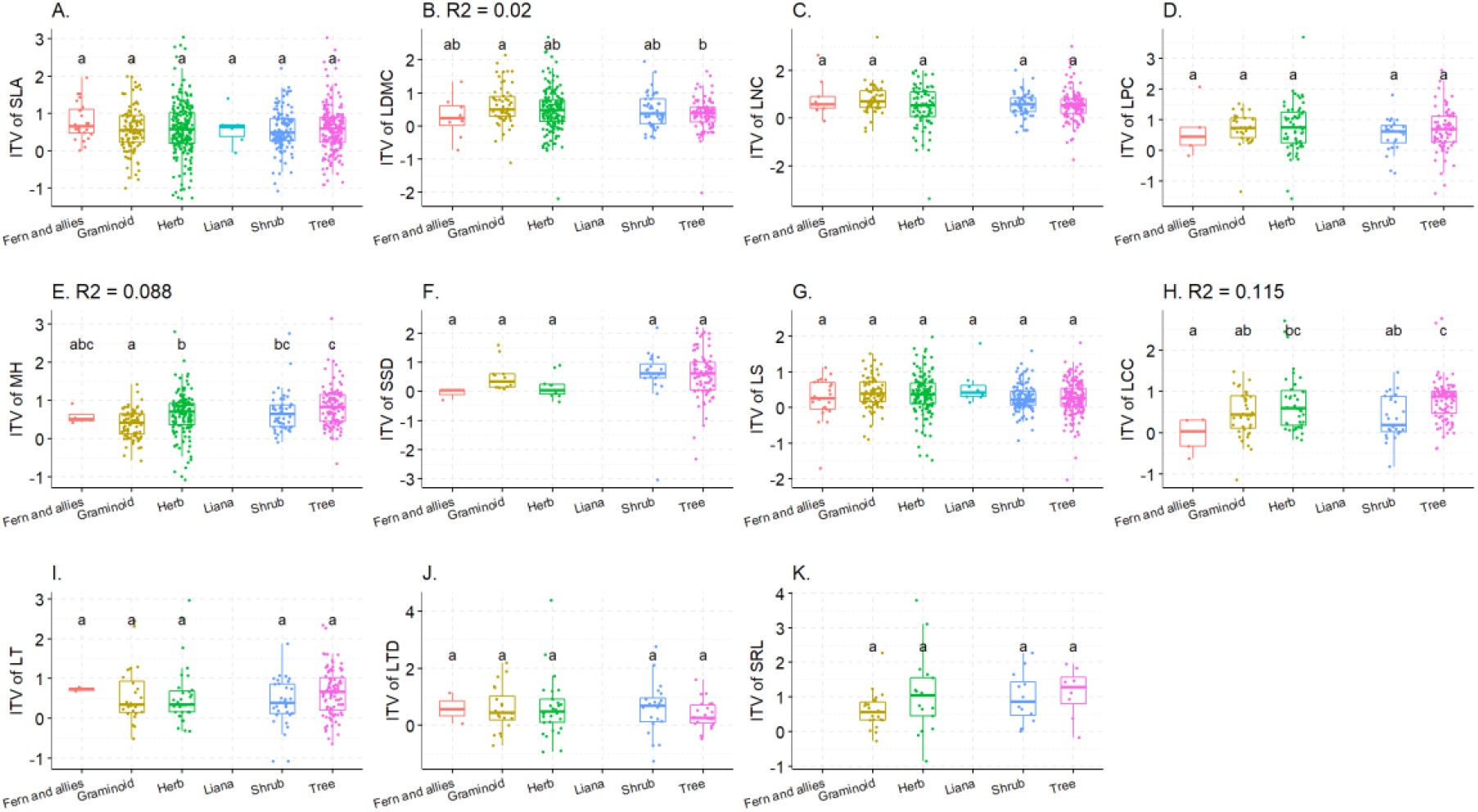
The intraspecific trait variation (ITV) of different traits vs. growth form (Fern and allies, Graminoid, Herb, Liana, Shrub, Tree), representing the influence of growth form on the ITV of A. specific leaf area, B. leaf dry matter content, C. leaf nitrogen content, D. leaf phosphorus content, E. maximum height and F. stem specific density, G. leaf size, H. leaf carbon content, I. leaf thickness, J. leaf tissue dense, K. specific root density, respectively. Different lowercase letters indicate that the means of ITV for a trait are significantly different (p < 0.05) between growth forms (as determined by Tukey post-hoc tests). R^2^ was only added when significant differences were detected.

### 3.3 Intraspecific trait variation and species alpha niche position

The ITVs of eight traits were related to either maximum or minimum species alpha niche position (Fig. **4**). The minimum species alpha niche position was negatively related to the ITVs of SLA (p < 0.001, R^2^ = 0.014) and SSD (p < 0.01, R^2^ = 0.067) while it was positively related to the ITVs of LPC (p < 0.01, R^2^ = 0.026), MH (p < 0.05, R^2^ = 0.017) and LCC (p < 0.01, R^2^ = 0.065). Moreover, the maximum species alpha niche position was positively related to the ITV of LDMC (p < 0.001, R^2^ = 0.121), leaf thickness (p < 0.001, R^2^ = 0.124) and leaf tissue density (p < 0.05, R2 = 0.046). There were no significant patterns for the other three traits.

**Fig. 4.**
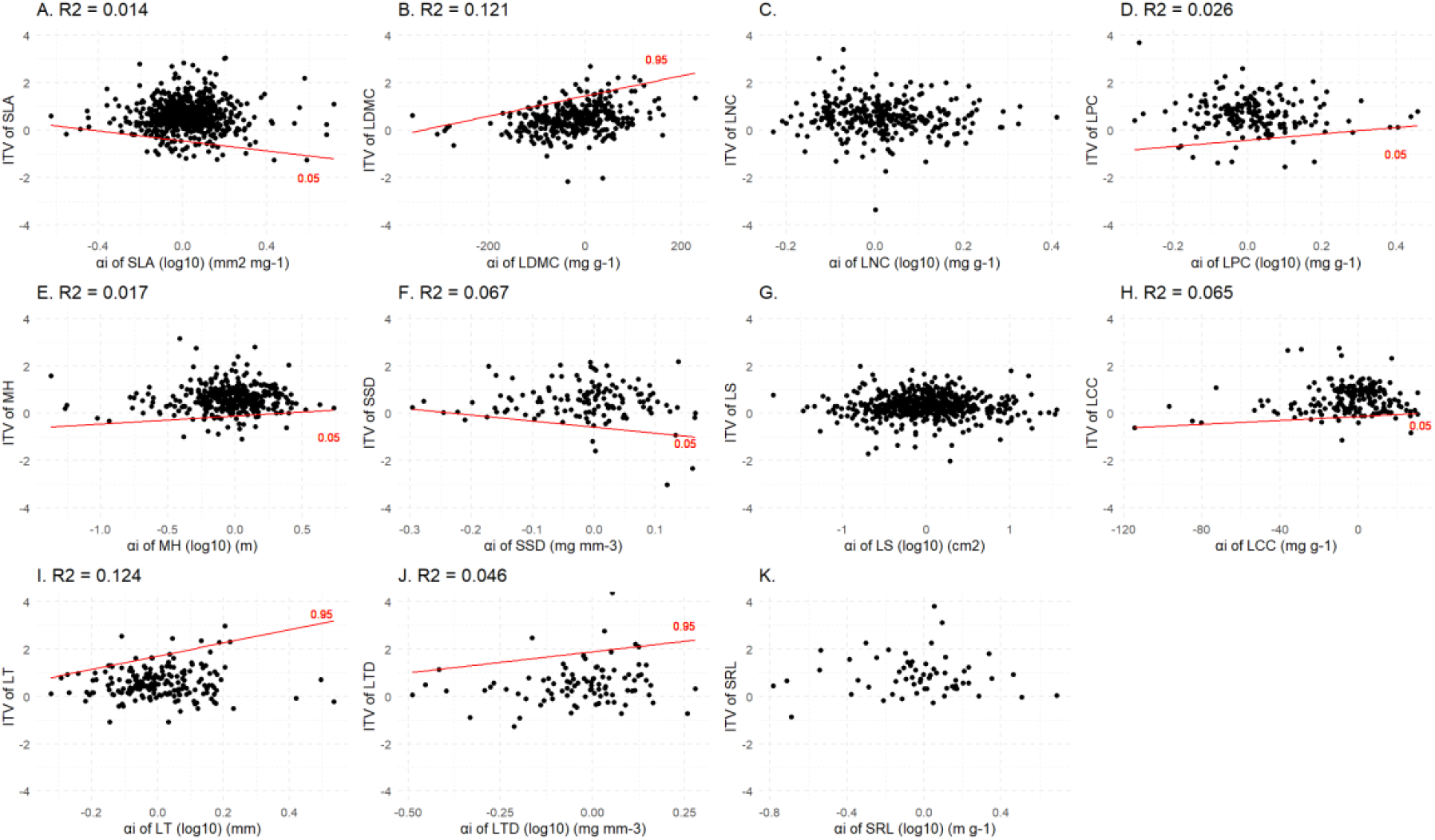
Quantile regression of intraspecific trait variation (ITV) of different traits vs. species alpha niche position (α_i_), representing the influence of maximum or minimum species alpha niche position on the ITV of A. specific leaf area, B. leaf dry matter content, C. leaf nitrogen content, D. leaf phosphorus content, E. maximum height, F. stem specific density, G. leaf size, H. leaf carbon content, I. leaf thickness, J. leaf tissue density, K. specific root length, respectively. The 0.05 and 0.95 quantile regression lines and R^2^ were only added when there were significant (p < 0.05) slopes.

### 3.4 Intraspecific trait variation and species beta niche position

The ITVs of seven traits were related to species beta niche position (Fig. **5**): LDMC, LNC, LPC, MH, SSD, LCC and SRL, while there was no significant pattern for the other four traits. Except for the ITV of LDMC which was negatively related to species beta niche position (p < 0.01, R^2^ = 0.023), species beta niche position always positively affected the ITV of LNC (p < 0.05, R^2^ = 0.018), LPC (p < 0.05, R^2^ = 0.032), MH (p < 0.01, R^2^ = 0.026), SSD (p < 0.05, R^2^ = 0.047) and SRL (p <0.05, R^2^ = 0.066). The positive relationship between species niche position and LCC (p < 0.001, R^2^ = 0.106) seemed to be driven by the pattern of woody species (Fig. **S5**).

**Fig. 5.**
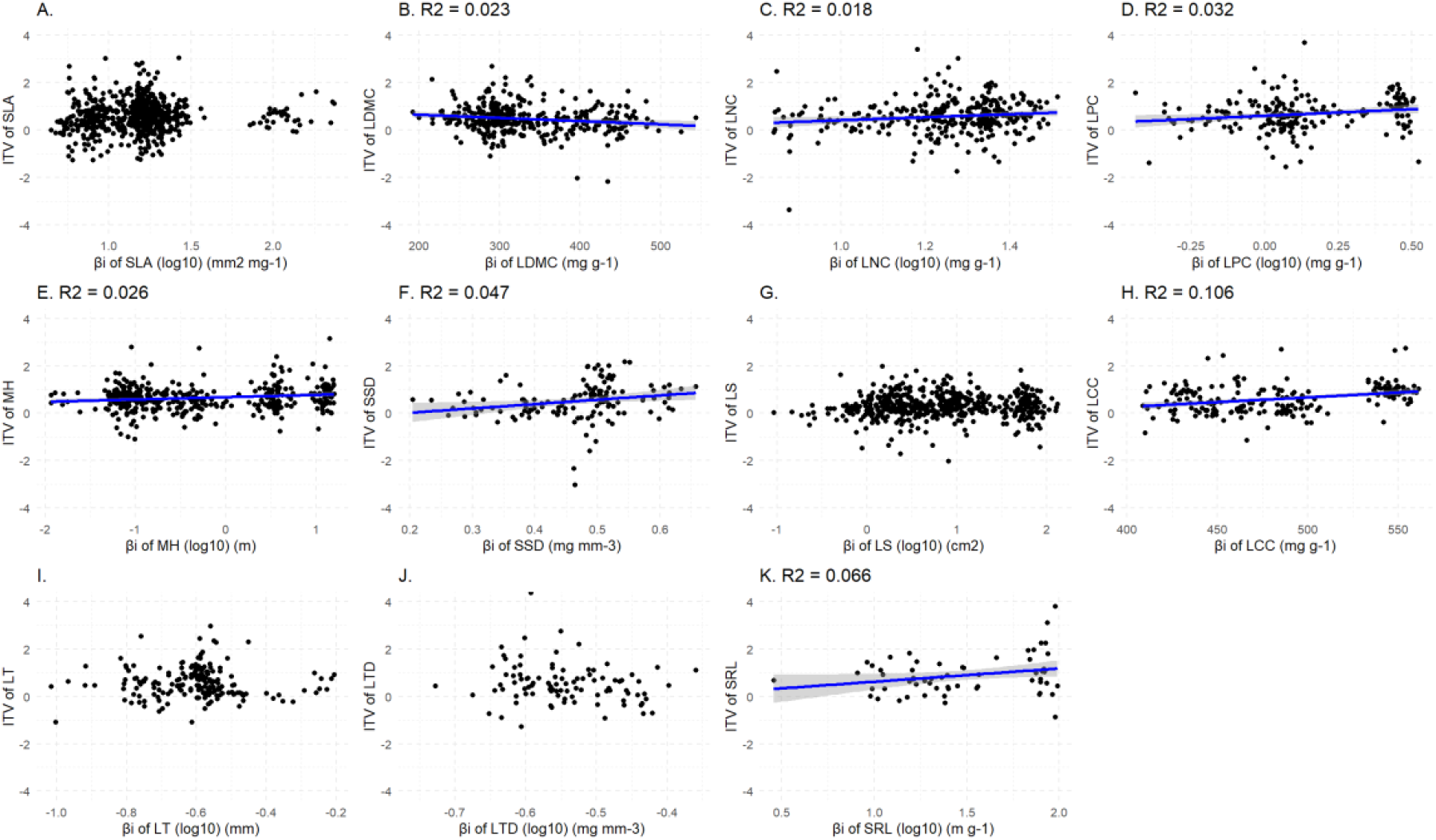
Ordinary least squares regression of intraspecific trait variation (ITV) of different traits vs. species beta niche position (β_i_), representing the influence of species beta niche position on the ITV of A. specific leaf area, B. leaf dry matter content, C. leaf nitrogen content, D. leaf phosphorus content, E. maximum height, F. stem specific density, G. leaf size, H. leaf carbon content, I. leaf thickness, J. leaf tissue density, K. specific root length, respectively. The regression lines and R^2^ were only added when there were significant (p < 0.05) slopes.

### 3.5 Intraspecific trait variation and species niche breadth

For the relationship between ITV and species niche breadth, only the ITV of LDMC was robustly positively related to species niche breadth (Fig. **6B**, p < 0.01, R^2^ = 0.263). For all other traits, the patterns were negligible or may be questionable due to the low number of observations (e.g. for LNC).

**Fig. 6.**
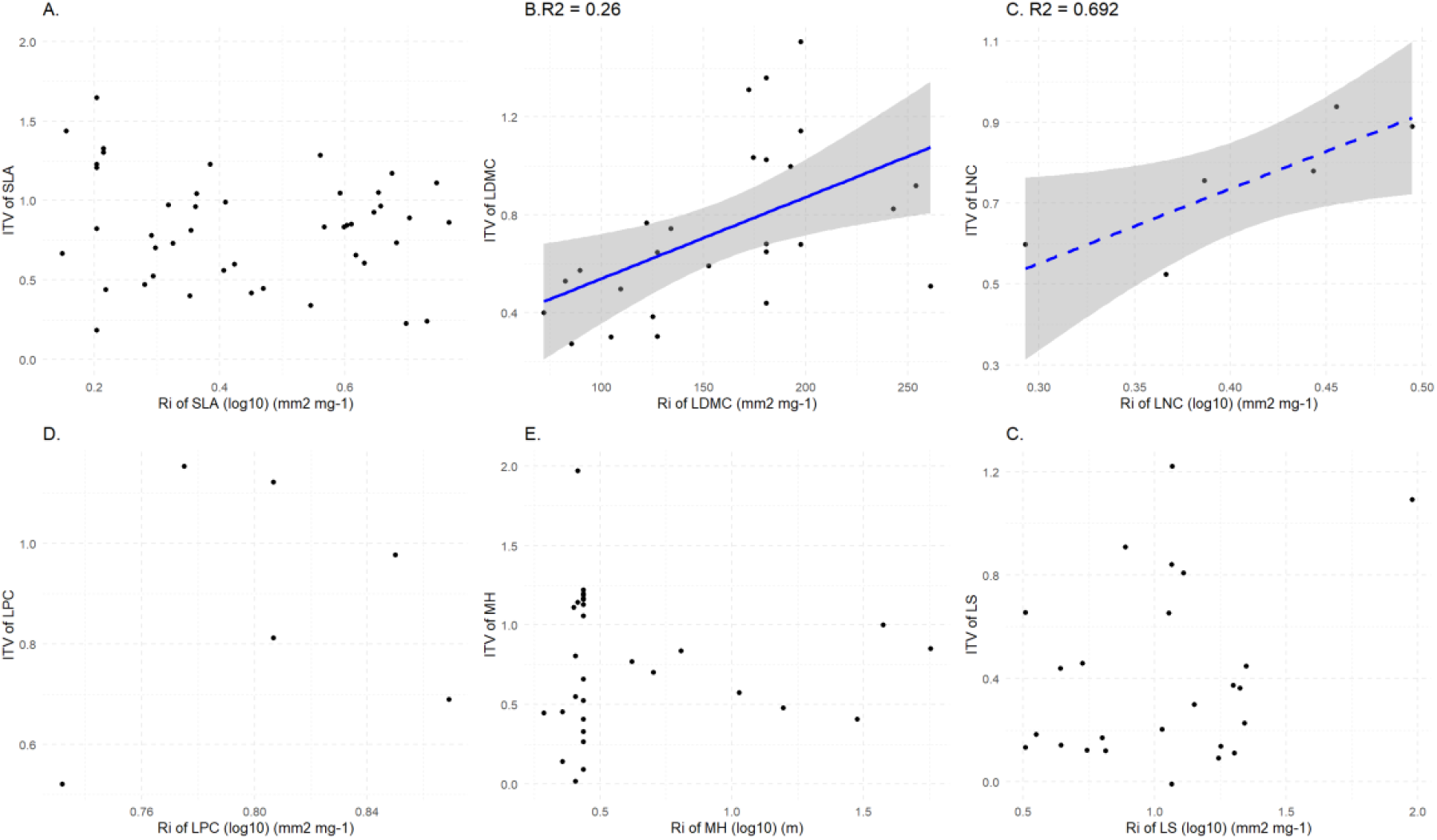
Relation between intraspecific trait variation (ITV) of different traits vs. species niche breadth (R_i_), representing the influence of species niche breadth on the ITV of A. specific leaf area, B. leaf dry matter content, C. leaf nitrogen content, D. leaf phosphorus content, E. maximum height, F. leaf size. Other traits are not shown as they did not have enough observations of niche breadth (n < 6) for this test. The regression lines and R^2^ were only added when there was a significant (p < 0.05) slope. The dashed regression line indicates the result of LNC may not be reliable as there are only 6 observations

### 3.6 Intraspecific trait variation and species CSR strategy

ITV could partly predict species CSR strategies for some traits (Table **1**). The ITV of leaf size was positively related to the percentage of species with a C strategy (p < 0.05, R^2^ = 0.008) and the percentage of species with a S strategy was negatively related to SSD (p < 0.05, R^2^ = 0.049), LCC (p < 0.05, R^2^ = 0.014) and leaf thickness (p<0.05, R^2^ = 0.037). There were no significant patterns for the other seven traits.

## 4 Discussion

### Lack of generic drivers of intraspecific trait variation

We aimed to explore the dominant and general drivers of intraspecific trait variation, by testing five species’ characteristics that are proxies for the way they cope with environmental conditions, biotic interactions and generic species features dealing with environmental and biotic conditions. Using a newly compiled database containing ITV estimates for 2064 species and 11 traits, our global-scale analysis provides the first comprehensive insight into the drivers of intraspecific trait variation. We found that environmental conditions, biotic interactions and species features all drove the ITV of some traits but those different traits had different main drivers. Therefore, none of these drivers can be considered a generic driver of ITV.

The absence of one dominantly generic driver of ITV may not be surprising in light of the multiple drivers that determine mean trait values on a global scale (Wright *et al*., 2004, 2017; Díaz *et al*., 2016). Regarding environmental conditions, we did account for the potential occurrence of multiple drivers: as a proxy of the expression of environmental conditions (and similar to Ackerly & Cornwell 2007), we assumed that the mean community trait value expressed the selection of this trait by multiple environmental drivers. If similar environmental drivers would also determine the variation in ITV among species, as assumed when designing this study, we would have found a strong relationship between the beta niche position and ITV. This was not the case. In contrast, the R^2^ of almost all relationships, including those ITV of traits more strongly related to biotic interactions, growth form and general species strategies, was very low. We, therefore, suggest that despite the comprehensive data compilation and analysis presented in this pioneering study, it is still early days for a global and collective understanding of the generic drivers of ITV.

### Drivers of intraspecific trait variation are highly trait-specific

Although individual traits showed idiosyncratic behaviour related to specific drivers, some trait-driver patterns - albeit with low explained variance - emerge from our analysis. In particular, the main proxy of environmental drivers, i.e. species beta niche position showed a relationship with the ITV of size-related traits (sensu Díaz *et al*. 2016) and leaf economics spectrum traits that reflect a plant’s resource use strategy (Wright *et al*., 2004; Shipley *et al*., 2006; Reich, 2014). Leaf morphological traits including SLA (although SLA is often also considered a leaf economics spectrum trait), in contrast, were hardly related to the species beta niche position. The latter is intriguing, as leaf morphological traits are very often also strongly related to environmental conditions, in particular, light and water availability, between species (Cornelissen *et al*., 2003; Pérez -Harguindeguy *et al*., 2013).

Our second proxy of environmental conditions, the species niche breadth, showed much weaker relationships with ITVs. This may be considered a bit unexpected as a large niche breadth would force species to adapt along a wide array of environmental conditions and vice versa a high ITV would allow for a large niche breadth. While the directionality of the causal relationship between ITV and niche breadth might be problematic, it is intriguing that only the ITV of LDMC showed a distinct relationship. This may be partly due to the substantially fewer number of observations for which the relationship with niche breadth could be tested.

In contrast to the relationships with environmental conditions, we found that biotic interactions – expressed through evaluating the relationships between ITV and species alpha niche position – were primarily significantly related to leaf morphology traits and size-related traits but less to leaf economics traits. This is consistent with a previous finding that the plasticity of some morphological traits (including SLA, leaf water content and maximum height) was related to the frequency with which neighbours were encountered (Abakumova *et al*., 2016). The 95^th^ percentile of ITV of several morphological traits (LDMC, leaf thickness and leaf tissue density) showed a positive relationship with the species alpha niche position. This indicates that species with maximum ITV of these traits tend to have higher trait values than other species within the community. These traits may correspond to species that are more stress tolerant, although a similar pattern was not found for the CSR strategy (Table **1**). For the 5^th^ percentile of the ITV, both negative and positive relationships with the species alpha niche position were found for some traits. The negative relationships for SLA and SSD indicate that the minimum feasible ITV value has to increase with more negative alpha niche positions. For example, when avoiding competition from species with higher SLA values, a higher value of ITV is needed. This may help the competing species to adapt to heterogeneous soil or microclimatic conditions. In addition to counter-gradient responses (Lusk *et al*., 2008; Liu *et al*., 2016), avoidance of competition may also explain why for SLA (and also other traits), negative ITVs were found for some species.

Since growth form and the CSR strategies are metrics that represent ways in which species deal with their environment and biotic interactions, we expected these to integrate a number of relevant drivers of ITV. However, compared to *α*_*i*_ and *β*_*i*_, there were fewer significant relationships with ITV for growth form and CSR strategies. Only MH and LCC showed significant and relatively strong patterns with growth form with the highest ITVs found in trees. This implies that in communities dominated by trees, likely coinciding with communities with major competition for light, there is a stronger demand for high variability in MH and LCC, possibly as a means to escape competition for light. These features may also explain the positive relationship with the beta niche position for these two traits and may shed some light on the relationship found for the alpha niche position for LCC and MH.

### Implications

The last few years have seen an increased interest in trait-based approaches for global modelling, a trend which we strongly support. However, the drivers of ITV seem strongly deviated from those of the species mean trait values. For example, we found growth form, species alpha niche position, species beta niche position and CSR were strongly related to species mean trait values (with variance explained R^2^ for *α*_*i*_ ranging between 0.06 - 0.71, and for *β*_*i*_ ranging between 0.25-0.92, see Fig. **S6-8** and Table **S1**), while these drivers hardly explained any variation in ITV (all R^2^ ≤ 0.12). These results suggest a decoupling of drivers affecting trait expression within species compared to those between species. Our observations corroborate the finding of Zhou *et al*. (2022) that some trait-trait relationships prevailing between species were not maintained in the ITV-ITV relationships of these traits. This decoupling has important implications for the use and understanding of trait-based approaches in general as well as may impact the way trait-based approaches are implemented in global modelling. The decoupling suggests that certain trait coordination may fall apart upon climate change, at least at the individual species level, but possibly also at the plant functional type level. This should be considered in global models for predicting global change impacts on vegetation dynamics. For example, most current Dynamic Global Vegetation Models (DGVMs) describe vegetation dynamics based on plant functional types (PFTs) and mostly ignore trait variation within PFTs (Moran *et al*., 2016). Implementing whether trait variation within PFTs is due to intraspecific or interspecific could improve those empirical approaches in modelling (e.g. van Bodegom *et al*. (2014) and Verheijen *et al*. (2013, 2015)) and evolutionary optimality approaches (Franklin *et al*., 2020). Moreover, models that derive or impose trait-trait relationships (e.g. Sakschewski *et al*., 2015)) should also take into account that certain trait coordination may fall apart. It is thus important not to hardwire trait-trait relationships such as leaf economic spectrum in global vegetation models. Instead, our findings provide a foundation for further refining these trait-based approaches to better account for alternative strategies.

While the current study does not provide definitive explanations on why some traits respond to certain drivers but not to others, we did find that some clusters of traits seem to have similar drivers of intraspecific trait variation. It warrants investigating why, for example, traits related to morphology show different patterns than the other traits.

Moreover, the low variance explained R^2^ associated with the drivers of ITV in this study highlights the lack of understanding of intraspecific trait variation to date. There have been suggestions that at within species level, species may develop different trait-trait relationships as alternative strategies (Umaña & Swenson, 2019) rather than respond to the same drivers by changing the same traits. It is also clear that genetic underpinnings may have an important bearing on ITV. Deciphering the respective roles of phenotypic plasticity and genetic differences in ITV might be an important next step in increasing our understanding of ITV. Such analysis, in combination with the analysis provided here, will fundamentally improve our understanding of the role of intraspecific trait variation in species eco-evolutionary strategies, as well as help predict vegetation responses to environmental change.

## Acknowledgements

J Zhou was funded by a PhD scholarship from China Scholarship Council (No. 201608310102). We thank the editor and anonymous reviewers for their constructive suggestions which significantly improved the manuscript.

## Author Contribution

P.M.v.B conceived the study; J.Z., E.C. and P.M.v.B developed the ideas; P.M.v.B and J.Z. collected the data; J.Z., E.C. and P.M.v.B conducted the analysis; J.Z. wrote the first draft. All authors contributed critically to the drafts and gave final approval for publication.

## Data Availability

The data that supports the findings of this study will be openly available in the Zenodo repository after acceptance.

## Supporting Information

**Fig. S1** The species niche breadth (R_i_) of log_10_-transformed specific leaf area (SLA) of species *Amomyrtus luma vs*. its observations (counts) for calculating its R_i_ from the bootstrap procedure.

**Fig. S2** Distribution of available species trait positions within C, S and R triangle in the database.

**Fig. S3** Species intraspecific trait variation (ITV) distribution of different traits.

**Fig. S4** Species intraspecific trait variation (ITV) of different traits.

**Fig. S5** Relationship between intraspecific trait variation (ITV) of leaf carbon content (LCC) and species beta niche position *β*_*i*_.

**Fig. S6** Ordinary least squares regression of species mean trait values *vs*. species beta niche position (*β*_*i*_) for different traits.

**Fig. S7** Ordinary least squares regression of species mean trait values *vs*. species alpha niche position (*α*_*i*_) for different traits.

**Fig. S8** Quantile regression of species mean trait values *vs*. species alpha niche position (*α*_*i*_) for different traits.

**Table S1** Overview of the results of statistic tests on the drivers of the different species mean trait values.

