## Supplemental Information for "Drivers of plant intraspecific variation are trait-specific"

Article title: Drivers of plant intraspecific variation are trait-specific

Article acceptance date: Click here to enter a date.

The following Supporting Information is available for this article:

**Fig. S1 The species niche breadth (**$\boldsymbol{R}_{\boldsymbol{i}}$**) of log10-transformed specific leaf area (SLA) of species *Amomyrtus luma* vs. its observations (counts) for calculating its** $\boldsymbol{R}_{\boldsymbol{i}}$ **from the bootstrap procedure** The black vertical line shows the mean of minimum number of observations (counts = 32) whose corresponding mean species niche breadth did not significantly differ from the maximum species niche breadth (calculated from 134 observations)

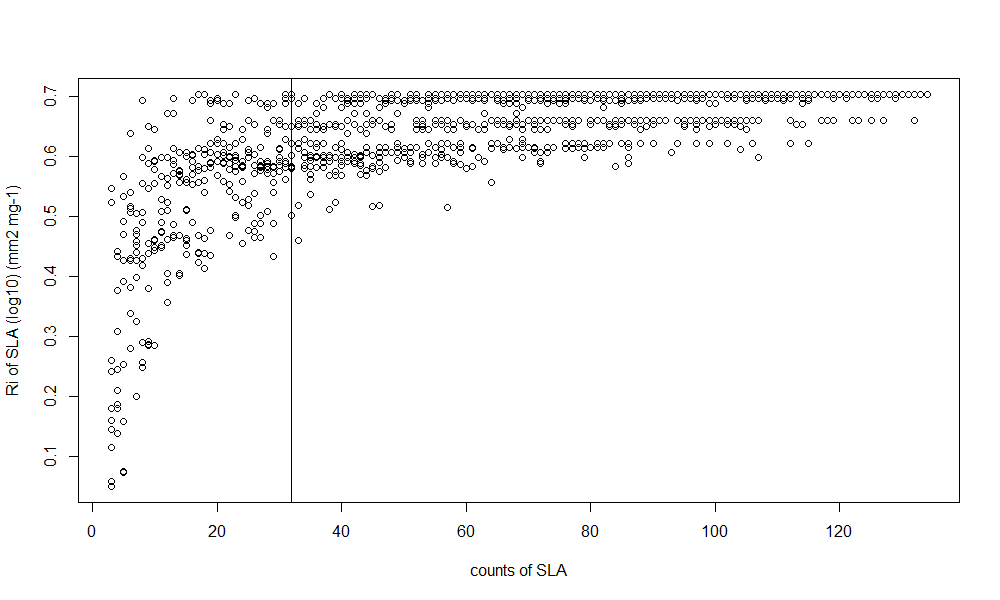

**Fig. S2** **Distribution of available species trait positions within C, S and R triangle in the database**.
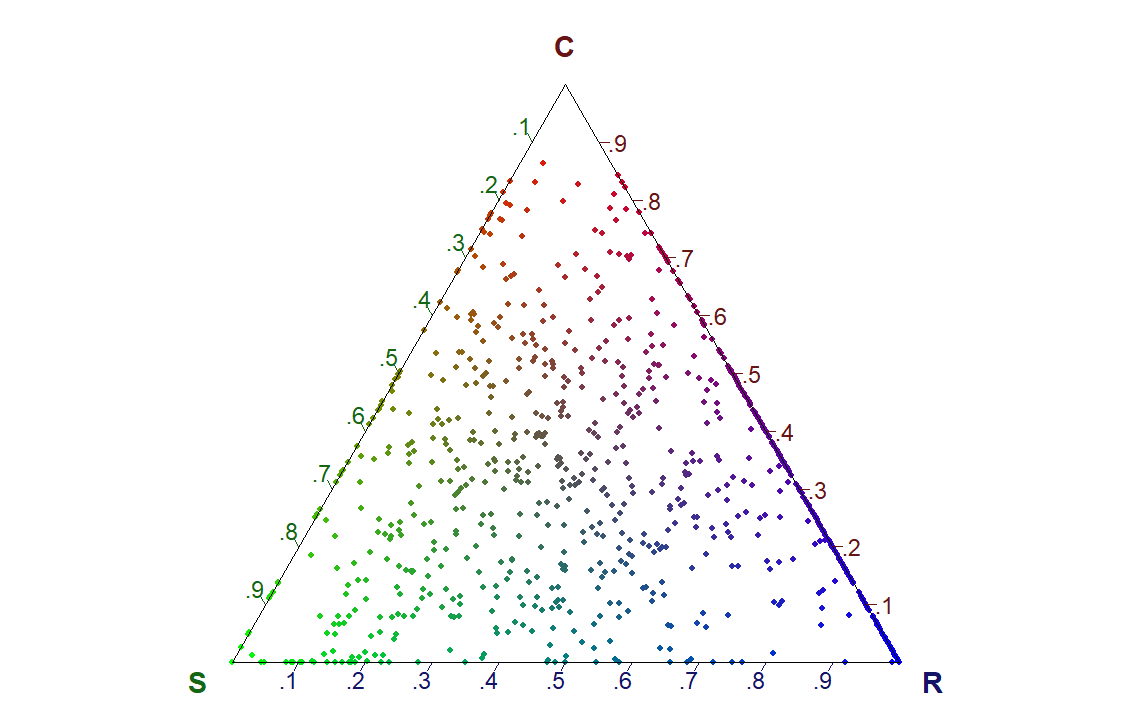

**Fig. S3 Species intraspecific trait variation (ITV) distribution of different traits**, representing the ITV distribution of different species for A. specific leaf area, B. leaf dry matter content, C. leaf nitrogen content, D. leaf phosphorus content, E. maximum height, F. specific stem density, G. leaf size, H. leaf carbon content, I. leaf thickness, J. leaf tissue density, K. specific root length, respectively. The black vertical line indicates the median values of ITV among species of each trait. Each square represents a species and different colours of the square indicate the number of observations of the species (i.e. the number of plots in which the species occurred) for estimating their ITV values.

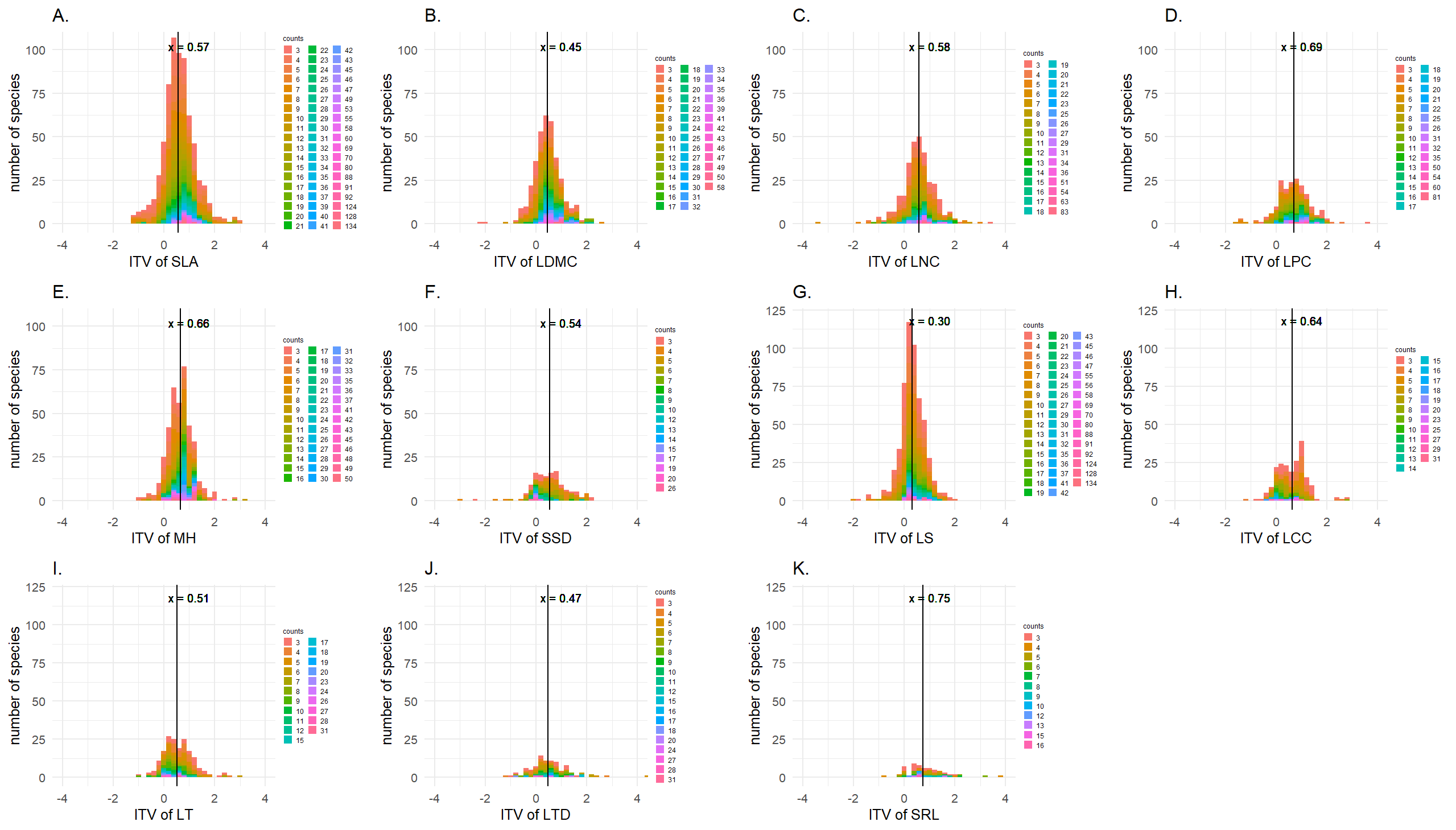

**Fig. S4 Species intraspecific trait variation (ITV) of different traits.** Leaf carbon content (LCC), leaf dry matter content (LDMC), leaf nitrogen content (LNC), leaf phosphorus content (LPC), leaf size (LS), leaf thickness (LT), leaf tissue density (LTD), maximum height (MH), specific leaf area (SLA), specific root length (SRL), stem specific density (SSD). Different lowercase letters indicate that the means of ITV for each trait are significantly different (p < 0.05) between each other (as determined by Tukey post-hoc tests).

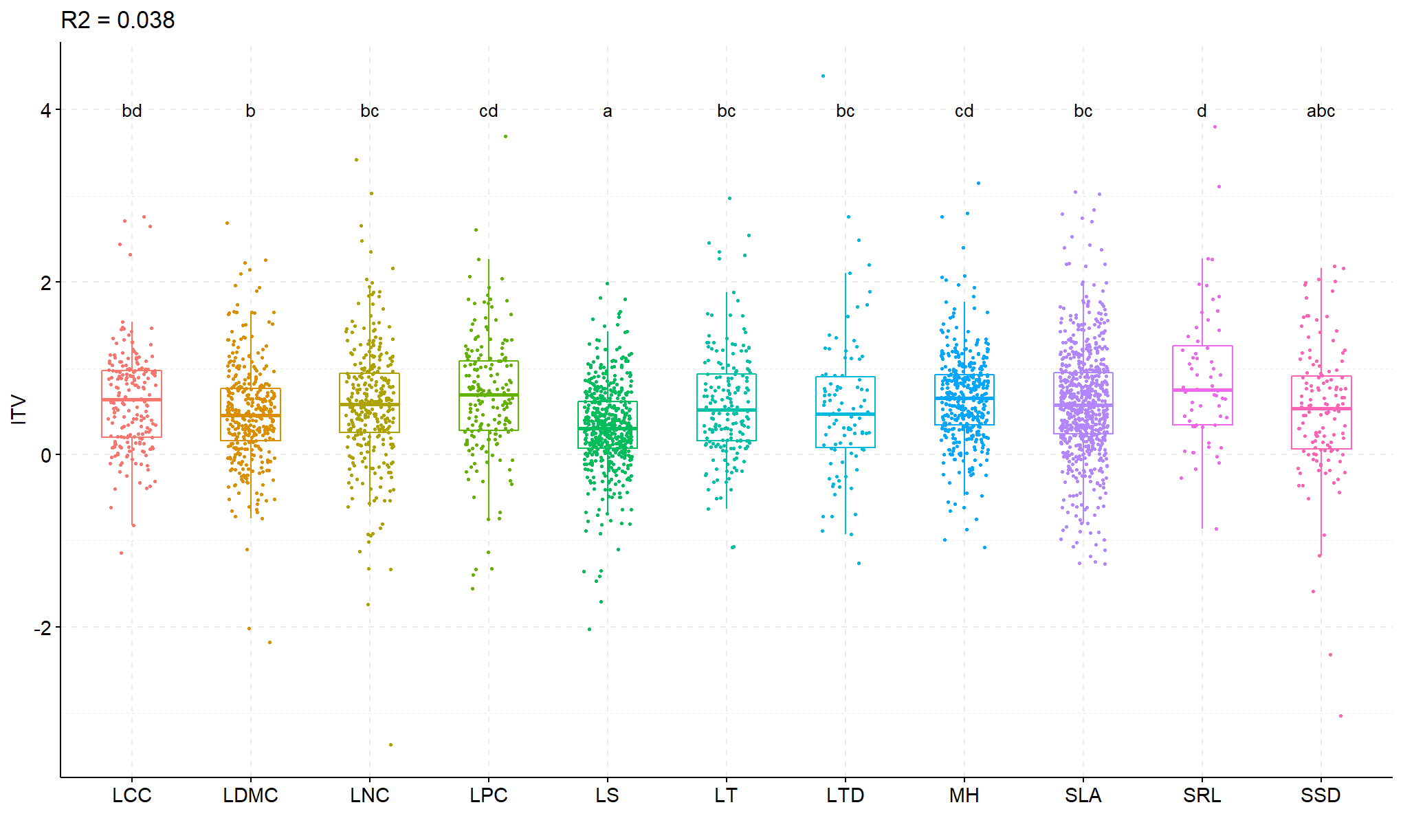

**Fig. S5** **Relationship between intraspecific trait variation (ITV) of leaf carbon content (LCC ) and species beta niche position** $\boldsymbol{\beta}_{\boldsymbol{i}}$**.** Non-woody species marked in red and woody species marked in blue.

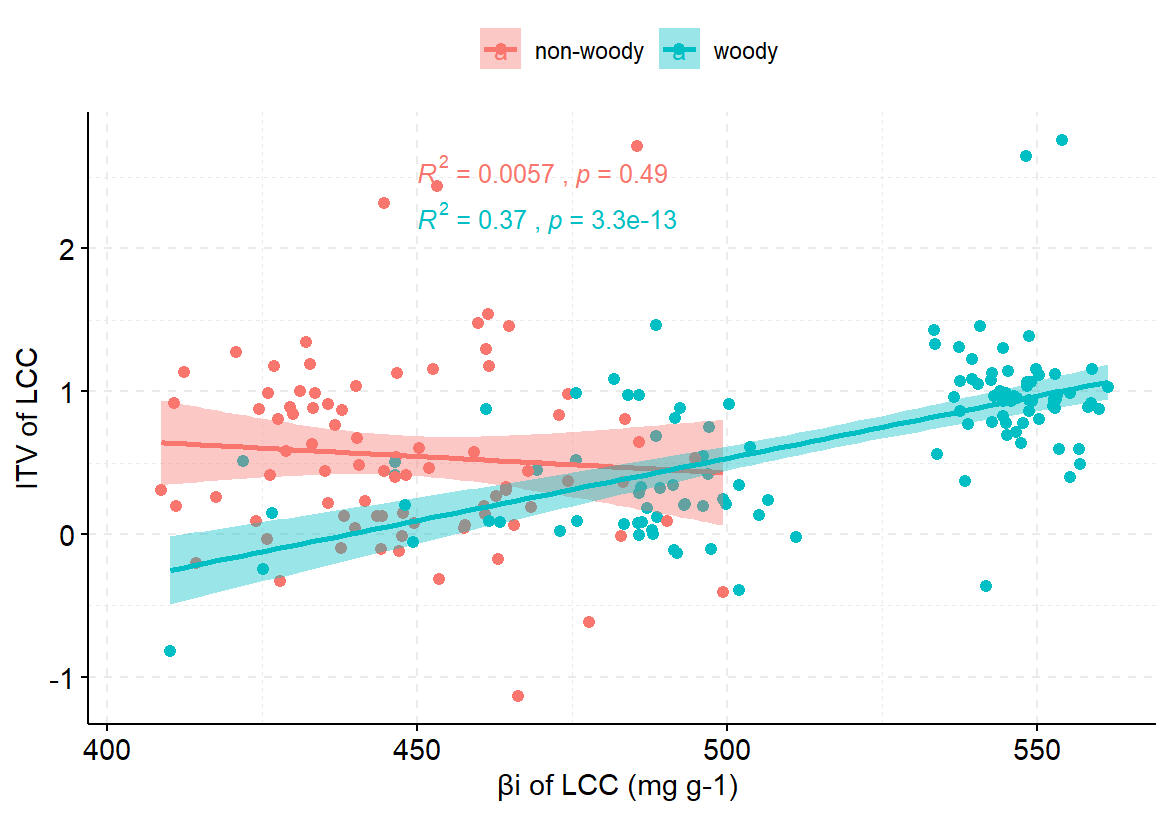

**Fig. S6 Ordinary least squares regression of species mean trait values *vs*. species beta niche position (**$\boldsymbol{\beta}_{\boldsymbol{i}}$**) for different traits** A. maximum height, B. specific stem density, C. leaf size, D. specific leaf area, E. leaf dry matter content, F. leaf carbon content, G. leaf nitrogen content, H. leaf phosphorus content, I. leaf thickness, J. specific root length, K. leaf tissue density. R^2^ and p-value are indicated for significant regressions.

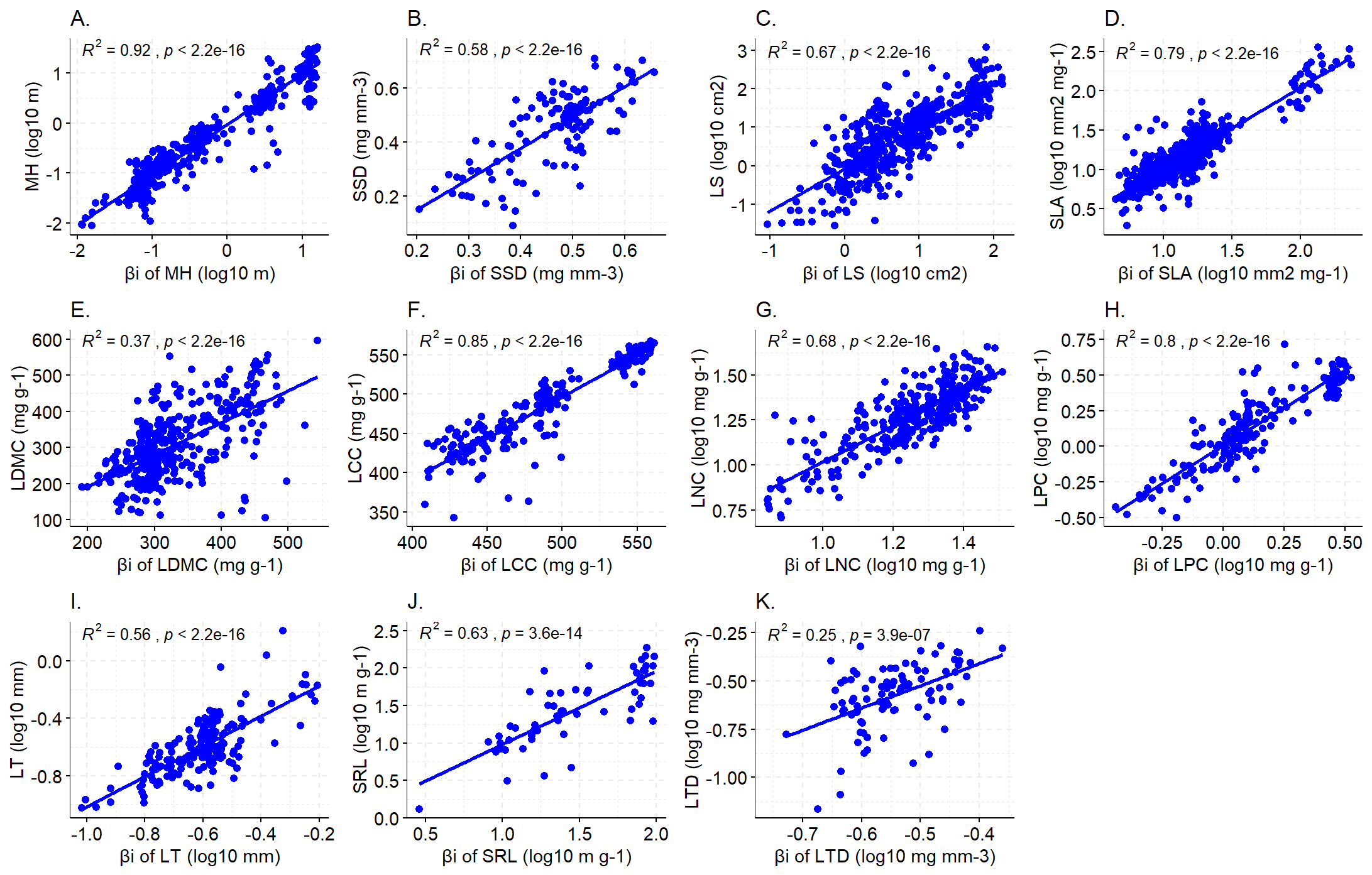

**Fig. S7 Ordinary least squares regression of species mean trait values *vs*. species alpha niche position (**$\boldsymbol{\alpha}_{\boldsymbol{i}}$**) for different traits.** A. maximum height, B. specific stem density, C. leaf size, D. specific leaf area, E. leaf dry matter content, F. leaf carbon content, G. leaf nitrogen content, H. leaf phosphorus content, I. leaf thickness, J. specific root length, K. leaf tissue density. R^2^ and p-value are indicated for significant regressions.

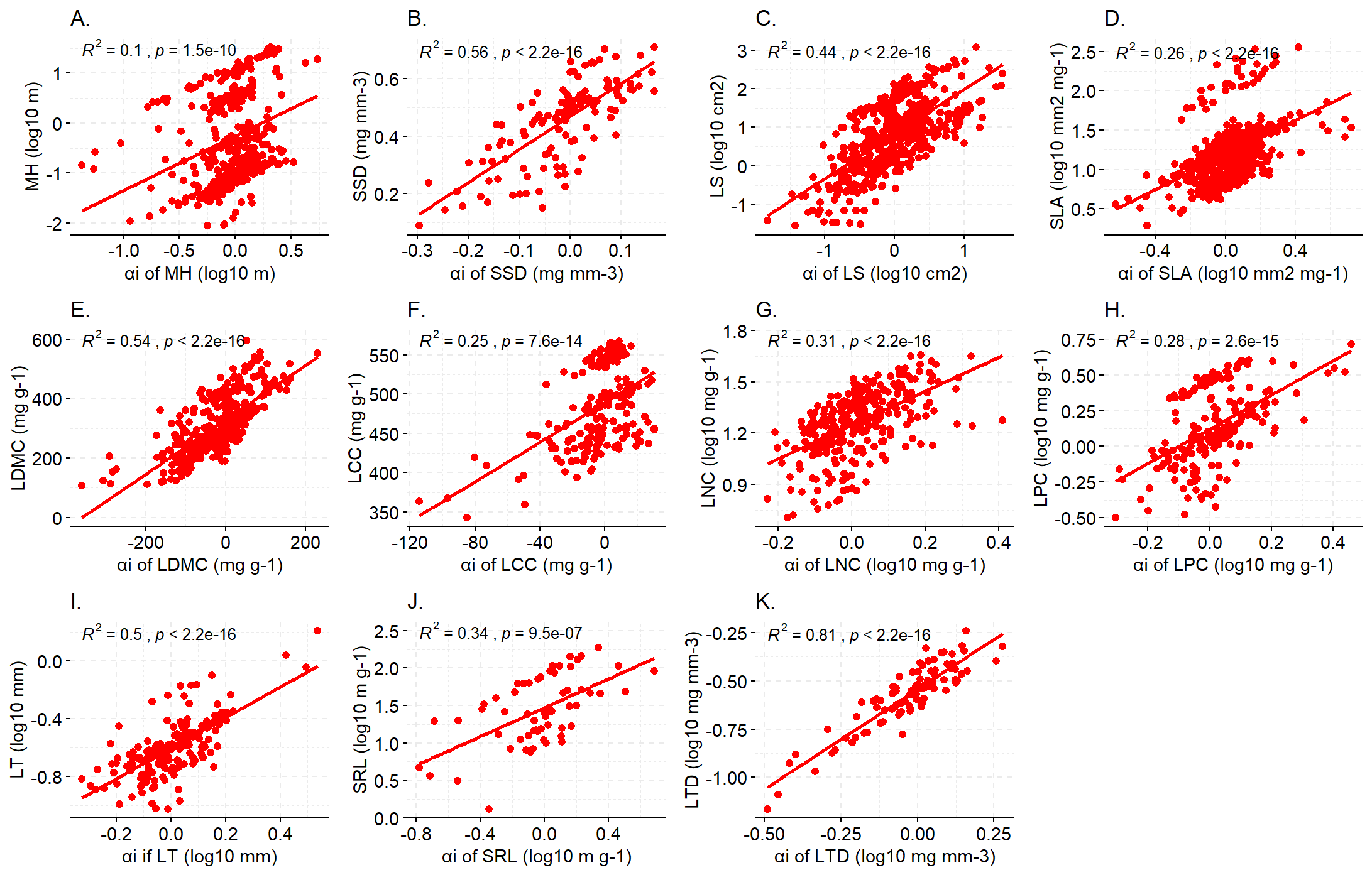

**Fig. S8 Quantile regression of species mean trait values *vs*. species alpha niche position (**$\boldsymbol{\alpha}_{\boldsymbol{i}}$**) for different traits.** A. maximum height, B. specific stem density, C. leaf size, D. specific leaf area, E. leaf dry matter content, F. leaf carbon content, G. leaf nitrogen content, H. leaf phosphorus content, I. leaf thickness, J. specific root length, K. leaf tissue density**.** Red lines indicate significant 0.95 quantile (above) and 0.05 quantile (below) regression lines.

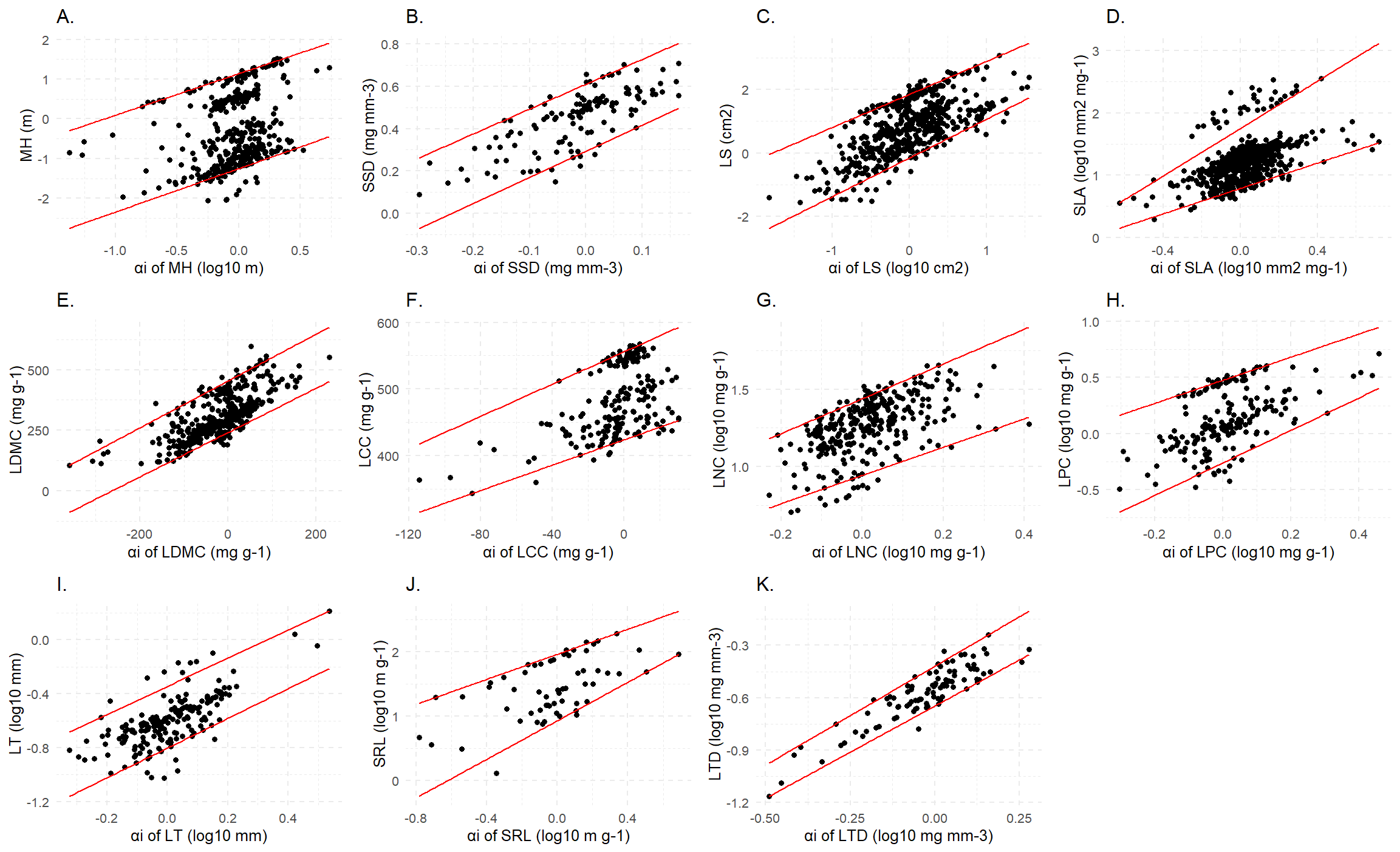

**Table S1** Overview of the results of statistic tests on the drivers of the different species mean trait values

| Trait  Driver | | | Leaf/root economics | | | | | Leaf morphology | | | | Size | |
| --- | --- | --- | --- | --- | --- | --- | --- | --- | --- | --- | --- | --- | --- |
|  |  |  | LNC | LPC | SRL | LCC | SLA | LDMC | LT | LTD | LS | MH | SSD |
|  | | # obs | 312 | 191 | 60 | 201 | 685 | 362 | 181 | 94 | 545 | 392 | 130 |
| Species feature | Growth form | R^2^ | 0.10 | 0.38 | 0.26 | 0.70 | 0.04 | 0.24 |  | 0.37 | 0.24 | 0.72 | 0.54 |
|  |  |  | F^ab^  G^ac^  H^a^  S^bc^  T^b^ | F^ab^  G^b^  H^c^  S^a^  T^a^ | G^b^  H^a^  S^a^  T^a^ | F^a^  G^b^  H^b^  S^c^  T^d^ | F^ac^  G^ab^  H^bc^  L^c^  S^bc^  T^a^ | F^ab^  G^b^  H^a^  S^b^  T^b^ | F^a^  G^a^  H^a^  S^a^  T^a^ | F^a^  G^c^  H^b^  S^c^  T^c^ | F^ab^  G^ab^  H^a^  L^abc^  S^b^  T^b^ | F^abc^  G^b^  H^a^  S^c^  T^d^ | F^a^  G^a^  H^a^  S^b^  T^b^ |
| Biotic | Min 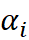 | R^2^ | 0.16 | 0.17 | 0.35 | 0.28 | 0.21 | 0.40 | 0.17 | 0.71 | 0.37 | 0.13 | 0.31 |
|  |  | Sig level | ***  (+) | ***  (+) | ***  (+) | ***  (+) | ***  (+) | ***  (+) | ***  (+) | ***  (+) | ***  (+) | ***  (+) | ***  (+) |
|  | (lm) 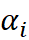 | R^2^ | 0.31 | 0.28 | 0.34 | 0.25 | 0.26 | 0.54 | 0.50 | 0.81 | 0.44 | 0.1 | 0.56 |
|  |  | Sig level | ***  (+) | ***  (+) | ***  (+) | ***  (+) | ***  (+) | ***  (+) | ***  (+) | ***  (+) | ***  (+) | ***  (+) | ***  (+) |
|  | Max 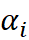 | R^2^ | 0.32 | 0.19 | 0.33 | 0.12 | 0.06 | 0.31 | 0.31 | 0.48 | 0.29 | 0.18 | 0.32 |
|  |  | Sig level | ***  (+) | ***  (+) | ***  (+) | ***  (+) | ***  (+) | ***  (+) | ***  (+) | ***  (+) | ***  (+) | ***  (+) | ***  (+) |
| Environment | 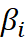 | R2 | 0.68 | 0.8 | 0.63 | 0.85 | 0.79 | 0.37 | 0.56 | 0.25 | 0.67 | 0.92 | 0.58 |
|  |  | Sig level | ***  (+) | ***  (+) | ***  (+) | ***  (+) | ***  (+) | ***  (+) | ***  (+) | ***  (+) | ***  (+) | ***  (+) | ***  (+) |
|  | $R_{i}$ | # obs | 6 | 6 | nd | nd | 45 | 26 | nd | nd | 25 | 30 | nd |
|  |  | R2 |  | 0.73 |  |  |  | 0.23 |  |  |  | 0.65 |  |
|  |  | Sig level |  | * |  |  |  | * |  |  |  | *** |  |
| Environment & Biotic | CSR | # obs | 191 | 119 | 35 | 148 | 354 | 277 | 95 | 79 | 288 | 233 | 66 |
|  |  | R^2^ | 0.05 | 0.02 | 0.14 | 0.05 | 0.14 | 0.08 | 0.10 | 0.09 | 0.17 | 0.02 | 0.10 |
|  |  | C | *** (+) | *  (+) |  |  | ***  (+) | *  (-) | ***  (-) |  | ***  (+) |  | ***  (-) |
|  |  | S | ***  (+) | **  (+) | *  (+) | **  (-) | ***  (+) | ***  (-) | ***  (-) | ***  (-) | ***  (+) |  | ***  (-) |
|  |  | R | *  (+) |  | *  (-) | ***  (-) | ***  (+) |  | **  (-) |  | *  (+) | *  (-) | ***  (-) |

ITV, intraspecific trait variation; Traits are indicated by their abbreviation: leaf nitrogen content (LNC), leaf phosphorus content (LPC), specific root length (SRL), leaf carbon content (LCC), specific leaf area (SLA), leaf dry matter content (LDMC), leaf thickness (LT), leaf tissue density (LTD), LS (leaf size), maximum height (MH), stem specific density (SSD); growth forms are indicated by their abbreviation: Fern and allies (F), Graminoid (G), Herb (H), Liana (L), Shrub (S), Tree (T); $\alpha_{i}$, species alpha niche position, $\beta_{i}$, species beta niche position, $R_{i}$, species niche breadth; “# obs” indicates the number of species for the test; different lowercase letters indicate that the means of species mean trait values are significantly different between different growth forms (as determined by Tukey post-hoc tests); “lm”, ordinary least squares regression; “nd”, no data; “***”, p < 0.001; “**”, p < 0.01; “*”, p < 0.05; “+”, positive relationship, “-”, negative relationship; results in grey may not be reliable as there are only 6 observations in the test; at the head of this table, the background colour of LCC and SLA are in orange indicating these traits are both leaf economics and leaf morphology traits, the background colour of LS is in light green indicating it is both leaf morphology and size-related traits.
